# Deep transcriptomic analysis of molecular mediators of complex traits in heterogeneous stock (HS) rats

**DOI:** 10.64898/2026.11.05.764420

**Authors:** Daniel B. Munro, Khai-Minh H. Nguyen, Celine L. St. Pierre, Hannah Bimschleger, Katarina A. Cohen, Yizhi Wang, Rachel Ward, Katie Holl, Denghui Chen, Benjamin B. Johnson, Gavrila Ang, Milad Mortazavi, Elaine Keung, Christopher King, Jordan A. Tripi, Brittany N. Kuhn, Brady M. Thompson, Arimantas Lionikas, Derek Daniels, Christopher Vulpe, Peter W. Kalivas, Olivier George, Douglas J. Adams, David M. Dietz, Shelly Flagel, Robert Hitzemann, Alexander Gusev, Laura M. Saba, Monica M. Jablonski, Thomas C. Jhou, Francesca Telese, Paul J. Meyer, Suzanne H. Mitchell, Leah C. Solberg Woods, Hao Chen, Oksana Polesskaya, Pejman Mohammadi, Abraham A. Palmer

## Abstract

**Background:** Rats are a powerful model organism for genome-wide association studies (GWAS) of physiological and behavioral traits. Heterogeneous Stock (HS) rats, which were created by intercrossing 8 inbred rat strains, are heavily used for this purpose because of their genetic diversity and accumulated recombinations due to over 100 generations of outbreeding, which improves mapping precision. However, functional genomics resources for post-GWAS interpretation of genomic findings have been limited.

**Results:** We processed 3,160 RNA-seq samples, 1,932 of which were previously unpublished, from 13 tissues as part of RatGTEx. We used Pantry to quantify six modalities of transcriptome regulation (expression, isoform ratio, intron excision ratio, alternative transcription start site (TSS), alternative poly(A), and RNA stability) in each sample, which allowed us to identify 144,876 independent quantitative trait loci (QTL) for 22,599 genes. We performed a transcriptome-wide association study (TWAS) with 19,796 unique genes across tissues and transcriptional modalities and 308 traits collected from 14 studies using FUSION, which identified 7,473 significant gene-trait associations.

**Conclusions:** This work presents a significant expansion of the RatGTEx resource in terms of samples, modes of transcriptome regulation, and genetic analyses. Our findings provide molecular evidence for the involvement of specific genes in mediating GWAS signals through a range of transcriptome regulatory mechanisms. These gene-to-phenotype relationships can be easily applied to genetic analyses in humans and other species. Further experiments can use this evidence to confirm these molecular mediators and target them for the treatment of behavioral and physiological traits and disorders. To facilitate use of these results, we have updated results on RatGTEx.org and provided a new interactive TWAS web portal (twas.ratgtex.org).

## Background

Outbred laboratory rats are commonly used to study the genetics of behavioral and physiological traits ^1–7^. Among the available rat populations, Heterogeneous Stock (HS) rats, which were created in 1984 by interbreeding eight inbred rat strains, have emerged as the population of choice because they provide both genetic diversity and accumulated recombinations that are necessary for fine-grained genetic mapping^8–10^. Although HS rats afford greater mapping precision than other available rat populations, fine mapping to individual variants, which is common in large human genetic studies, is limited in HS rats due to extensive linkage disequilibrium (LD) stemming from the eight inbred founders. Once genome-wide association study (GWAS) loci are found, the next step is typically to attempt to implicate genes that mediate the genetic associations. Sometimes one or more associated variants affect the protein sequence; however, based on findings in humans, most causal variants are presumed to be in non-coding regions and therefore to act through transcriptional regulation^11^.

Thus, RNA-seq data is now frequently collected to investigate the molecular underpinnings of complex trait genetic associations. These data are most commonly used to quantify gene expression, but genetic variants can affect many forms of variation such as different types of alternative splicing and transcription start and end sites. It is therefore important to quantify all of these transcriptional modalities, and tools such as Pantry streamline this process^12^.

Transcriptome regulatory models can also be used to predict gene expression and other transcriptional phenotypes for use in genetic studies to examine the role of molecular mediators. For example, after observing gene expression differences associated with self-administration of cocaine, expression predictions were used to distinguish between genetically regulated expression in genes that potentially mediate the amount of cocaine consumed, and expression differences resulting only from differential cocaine exposure^13^. Predictions can also be useful in studies where transcriptomic profiling is not performed; when a gene of interest has sufficiently heritable expression, its predicted expression can be analyzed alongside experimental data to test gene-specific hypotheses^14^. Another application is transcriptome-wide association study (TWAS), which integrates predictive transcriptomic models with GWAS results to identify potential mediating genes^15–17^.

The Rat Genotype-Tissue Expression (RatGTEx) project was introduced in 2022, providing quantitative trait loci (QTLs) affecting gene expression (eQTLs) and splicing (sQTLs) for five brain regions^18^ in *Rattus norvegicus*. Since then, the resource has greatly expanded in terms of number of brain regions and other tissues included, number of samples per tissue, breadth of transcriptomic variation assayed, and integration with GWAS data.

In this study, we report the largest collection of transcriptomic data in outbred rats and several new analyses of these data. First, we used Pantry^12^ to measure six modalities of transcriptomic variation for all samples. Next we mapped molecular QTLs (xQTLs) and created predictive models of gene expression and the other transcriptional modalities for all tissues. We then used these transcriptomic models to apply TWAS to a large collection of GWAS data from behavioral and physiological studies. We demonstrate that the resulting associations provide evidence for many mediators, including their specific modes of regulation, of the genetic effects on substance-use-related and other complex traits in rats.

## Results

We present an integrative analysis of genome-wide association study (GWAS) signals of behavioral and physiological traits in outbred rats with genetic drivers of transcriptome regulation (**Figure 1**). First, we collected RNA-seq data from 13 tissues (9 brain regions, whole brain hemisphere, liver, visceral adipose and eye tissue) from eight cohorts of genotyped heterogeneous stock (HS) rats (**Supplementary Table S1**). Then, in the latest phase of the RatGTEx project, we processed these data to quantify transcriptional phenotypes across six modalities, mapped cis-regulatory genetic associations, and built predictive transcriptomic models. Finally, we assembled GWAS data for 308 complex behavioral and physiological traits from 14 studies (**Supplementary Table S2**), applied multimodal transcriptome-wide association study (xTWAS), and analyzed the resulting putative transcriptional mediators.

**Figure 1.**
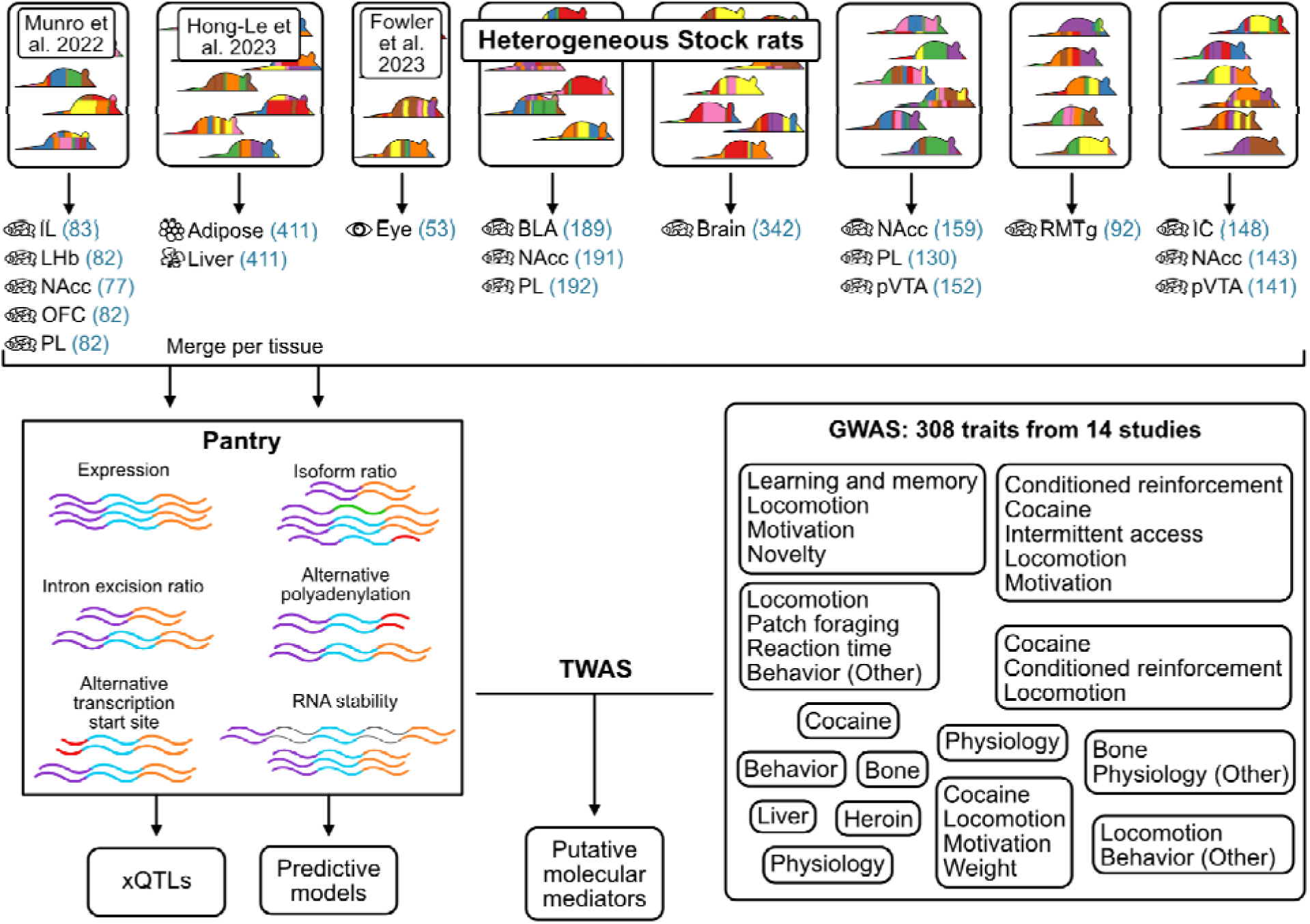
Overview of RatGTEx and Rat TWAS Hub. The diagram shows HS rat cohorts, the tissues extracted from each cohort, the transcriptomic data generated from them resulting in xQTLs and predictive models, and integration with external GWAS data to identify putative molecular mediators. HS rat cohorts are ordered by date added to RatGTEx, with Munro et al. 2022 being the pilot RatGTEx publication and additional datasets being published independently. Numbers in parentheses are the sample sizes obtained for each tissue after quality control filtering. Full tissue names are given in Supplementary Table S1. Each box within the GWAS box indicates one study and the categories of traits it provides.

We performed quality control steps to ensure correct matching of RNA-seq samples to genotypes. Of the 3,160 post-quality-control tissue samples, 71% underwent Illumina paired- end RNA-seq (read lengths 100 to 150), and the rest used single-end RNA-seq (read lengths 75 to 100). Mean reads per sample varied from 26.6 million to 41.4 million across datasets.

Using Pantry^12^, we quantified six modalities of transcriptional variation across each tissue dataset: gene expression levels, isoform ratio, junction-level intron excision ratio, alternative transcription start site (TSS) usage, alternative polyadenylation (poly(A)) site usage, and RNA stability. Hereafter, we refer to these quantified variables collectively as transcriptional phenotypes and to their six categories as transcriptional modalities. Whereas gene expression always produces only one phenotype per gene, several of the other modalities can produce multiple per gene. In total, we generated 129,777 to 182,667 (median 162,464) transcriptional phenotypes per tissue. The modality with the fewest phenotypes was RNA stability (10,284 to 13,901), whereas intron excision ratio had the most (32,809 to 58,092) (**Figure 2a**). More genes had expression phenotypes (median 24,071) than other modalities of phenotypes (medians ranging from 5,083 to 11,891, **Figure 2b**).

**Figure 2.**
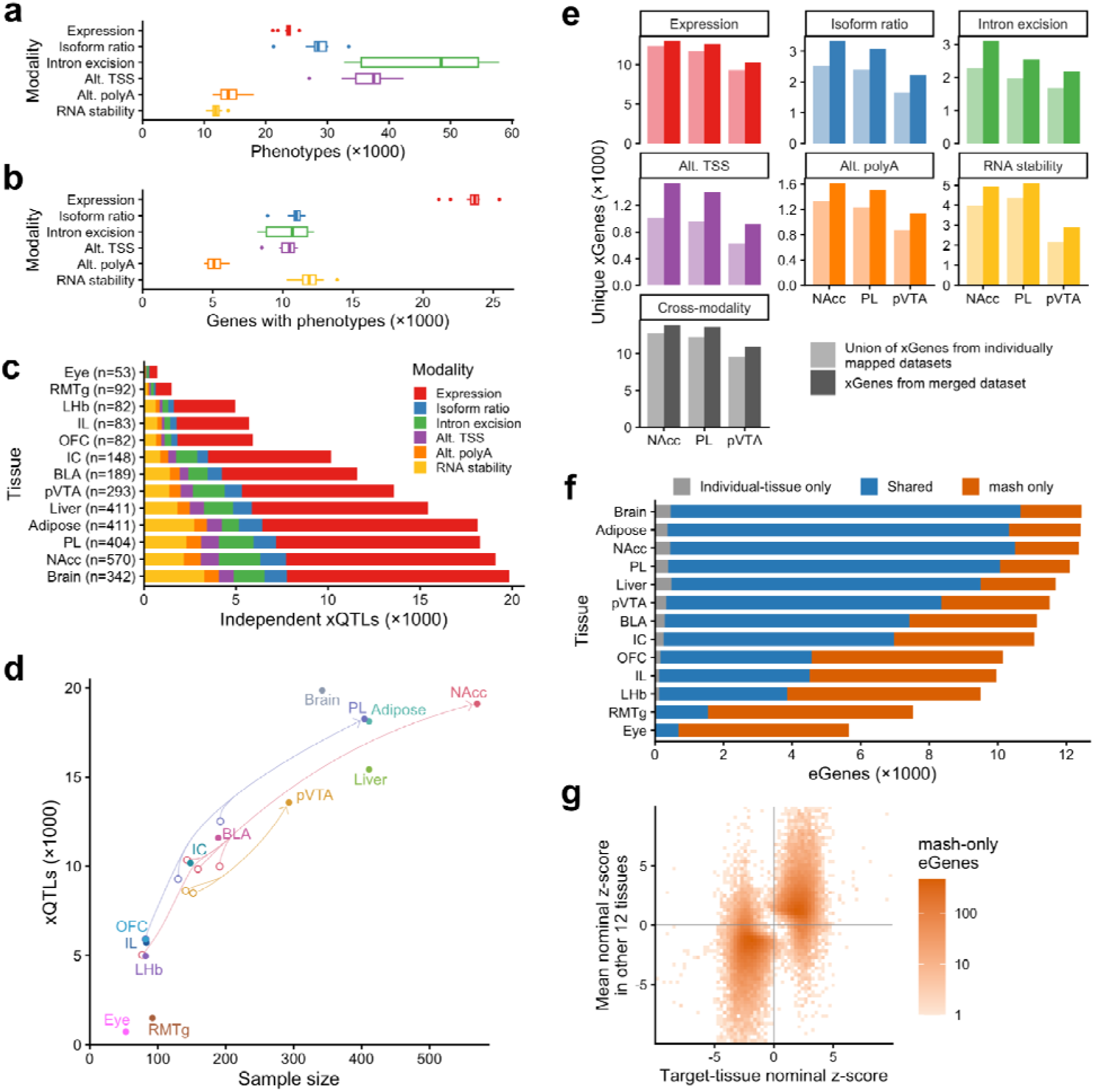
Multimodal phenotypes and xQTLs. **a** Boxplots show the number of transcriptional phenotypes of each modality generated for each tissue. **b** Boxplots show the number of genes with one or more phenotypes of each modality generated for each tissue. **c** Number of cis-xQTLs mapped for each tissue, colored by the modality of the mapped phenotype. Cross-modality mapping was performed, so xQTLs for the same gene in the same tissue are conditionally independent across modalities. **d** Number of independent xQTLs versus sample size per tissue. For each tissue with samples from multiple cohorts, the number of xQTLs and samples for each pre-merged sample set and for the merged set are both shown, connected by arrows. **e** Bar plots show the number of unique genes with xQTLs when mapping each pre-merged dataset individually (and taking the union of genes) compared to the xQTL genes when mapping the merged dataset. There were four, three, and two datasets for the NAcc, PL, and pVTA tissues, respectively. **f** eGenes from individual-tissue eQTL mapping and cross-tissue multivariate adaptive shrinkage (mash), colored by whether they were eGenes for individual-tissue mapping, mash, or both. **g** 2D histogram showing, for each mash-only eGene in **f**, the individual-tissue mapping nominal z-score for the top gene-variant pair in the eGene tissue versus the mean nominal z-score across the other 12 tissues. Axis limits were set to symmetrically include 99.5% of the data points.

### xQTL mapping

We mapped cis-QTLs for all transcriptional modalities for all expressed protein-coding and long noncoding RNA (lncRNA) genes in each tissue using Pantry’s cross-modality mapping, which uses tensorQTL^19^ to map xQTLs that are conditionally independent across all modalities per gene. We found 723 to 19,852 (median 11,587) independent xQTLs per tissue, affecting 3% to 61% (median 43%) of expressed genes, with similar proportions of modalities represented in each tissue (**Figure 2c**, **Supplementary Table S3**). The number of xQTLs per tissue is largely explained by sample size (**Figure 2d**), while differences in RNA-seq quality and depth could also affect mapping power (**Supplementary Figure S1**). For three tissues, we had samples from multiple cohorts–four for nucleus accumbens core (NAcc), three for prelimbic cortex (PL), and two for posterior ventral tegmental area (pVTA) (**Figure 1**). Mapping these as one merged dataset per tissue resulted in substantially more xQTLs than when mapping any constituent dataset separately (**Figure 2d**). In fact, for each tissue and for every modality and for cross-modality mapping, the number of unique genes with xQTLs mapped from the merged dataset exceeded the size of the union of genes with xQTLs mapped from individual datasets (**Figure 2e**). Batch effects resulting from the merging of cohorts prepared and sequenced separately were prominent in the normalized phenotypes but effectively captured by the phenotype principal component (PC) covariates (**Supplementary Figure S2**).

These results, corresponding to the RatGTEx v4 release, include 11.5-fold more xQTLs (when mapping each modality separately) than the pilot study^18^, due to new tissues, new samples for some tissues, and more transcriptional modalities. But even for the same raw input data, developments to the data processing pipeline have resulted in more independent eQTLs than previous releases (releases 1-3 do not have associated publications). For the five tissue datasets reported in the pilot study (infralimbic cortex (IL), lateral habenula (LHb), NAcc, orbitofrontal cortex (OFC), and PL from 86 rats), we found an average of 43% more eQTLs with the current pipeline, even before merging the additional NAcc and PL samples from newer cohorts.

To assess the ability to combine information across tissues to increase discovery power, we applied multivariate adaptive shrinkage (mash)^20^ to the eQTL summary statistics for all 13 tissues (**Supplementary Table S4**). This resulted in 1,784 to 5,972 additional genes with significant eQTLs (eGenes) per tissue, with the larger increases occurring in tissues with the fewest individual-tissue eGenes (**Figure 2f**). To see whether the mash-only eGenes were generally plausible as associations with signal boosted by cross-tissue information, we compared each top gene-variant pair’s nominal (single-tissue) z-score to the same pair’s mean nominal z-score across the other 12 tissues, observing concordant signs for 90.8% of them (**Figure 2g**).

### Predicting regulatory variation from genotype data

We used FUSION^16^ to build prediction models of gene regulation using genotype data and the transcriptional phenotypes for all modalities in each tissue. These models use a selection of variants within 500 kb of each gene’s TSS to predict transcriptional phenotypes for a genotyped individual. Gene expression was the most common modality among heritable phenotypes (P<0.01), with a median across tissues of 7,126 genes with predictions. A median across tissues of 1,731 additional genes had other-modality predictions only, while a median of 2,814 genes had both expression and other-modality predictions (**Figure 3a**). A median of 61% of the genes with predictions for a modality had such predictions in more than one tissue (**Figure 3b**).

**Figure 3.**
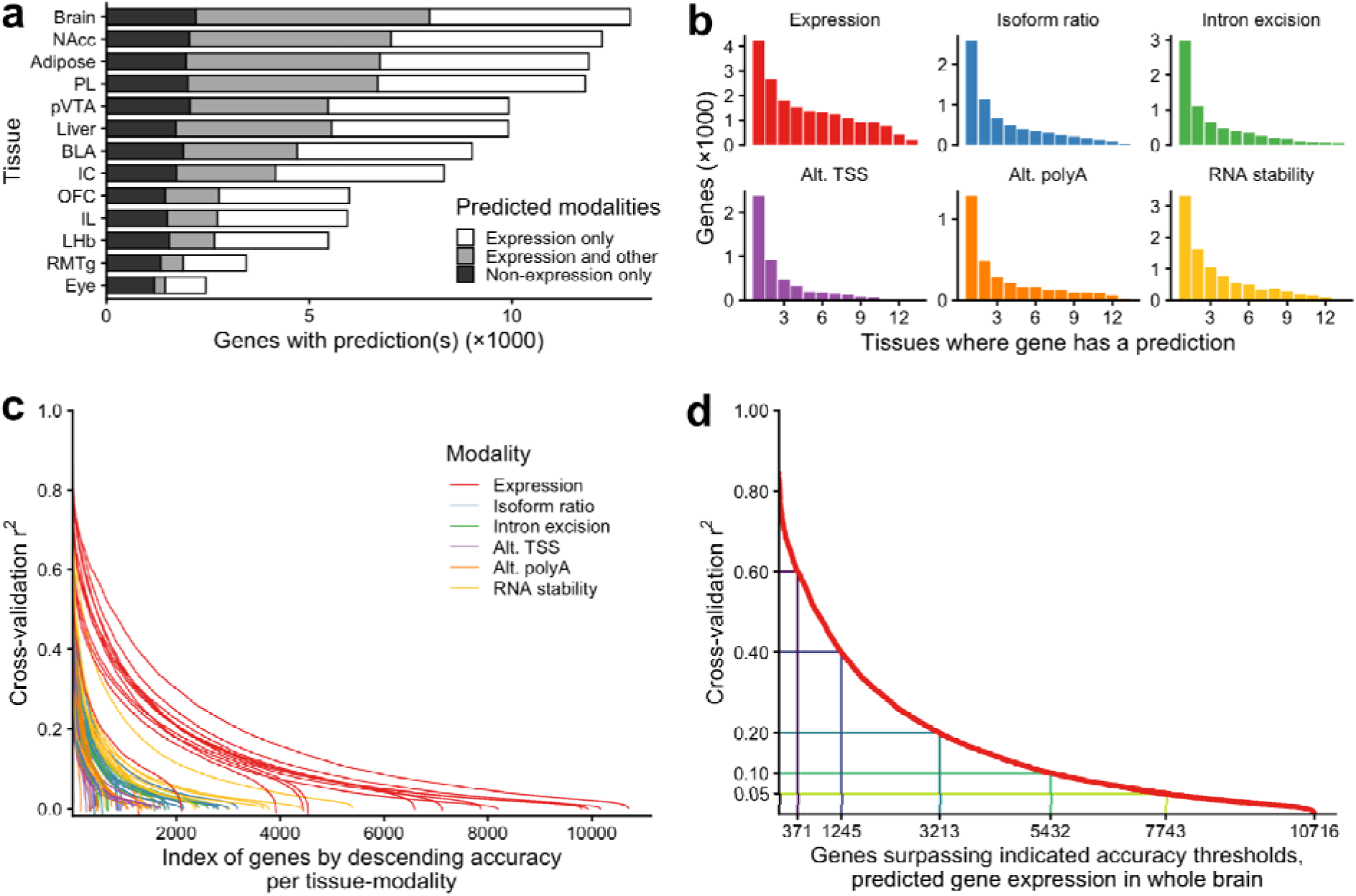
Transcriptional phenotype predictions. **a** Number of genes per tissue with transcriptional phenotype predictions, colored by whether they have predictions for expression, only other transcriptional phenotypes, or both. **b** For the genes with phenotype predictions of each modality, bars show the number of tissues for which the gene has a predicted phenotype. **c** Each curve shows prediction accuracy for one modality in one tissue. For each curve, genes are ordered on the x-axis by descending cross-validation adjusted r^2^, or mean r^2^ for genes with multiple predicted phenotypes of the same modality. **d** The curve for expression in whole brain reproduced from **c**, with colored lines indicating the number of genes exceeding various r^2^ thresholds.

Consistent with its greater heritability, expression could typically be predicted more accurately and for more genes than the other modalities (**Figure 3c, Supplementary Figure S3**). In whole brain, which produced the most expression predictions (10,716), there were 5,432 genes with cross-validation adjusted r^2^ of at least 0.1, 3,213 genes with r^2^ >= 0.2, and 1,245 genes with r^2^ >= 0.4 (**Figure 3d**).

### xTWAS

To study the transcriptomic mediation of GWAS associations and link genes to complex traits, we used the transcriptional models to run TWAS against traits from HS rat GWAS studies. We ran TWAS on 308 complex traits (**Supplementary Table S5**) using FUSION, generalized to multiple transcriptional modalities. Across all tissues, there were 7,473 unique gene-trait pairs among 50,137 total significant associations (**Figure 4a**, **Supplementary Table S6**). 28% of these pairs did not include hits involving the expression level modality in any tissue and were found only by including the other transcriptional modalities, highlighting the value of our approach. The total number of xTWAS hits per trait across tissues and modalities correlates with the trait’s GWAS sample size with Spearman ρ = 0.26 (**Figure 4b**). Transcriptional phenotypes with higher prediction accuracy tended to produce xTWAS hits with higher rates of genetic colocalization evidence: among significant xTWAS hits, the fraction with colocalization probability >0.8 increased from 1.2% for phenotypes with cross-validation r^2^ ≤ 0.05 to 7.9% for those with r^2^ > 0.4 (**Supplementary Figure S4**). Many loci with xTWAS hits included traits from multiple categories (**Supplementary Figure S5**).

**Figure 4.**
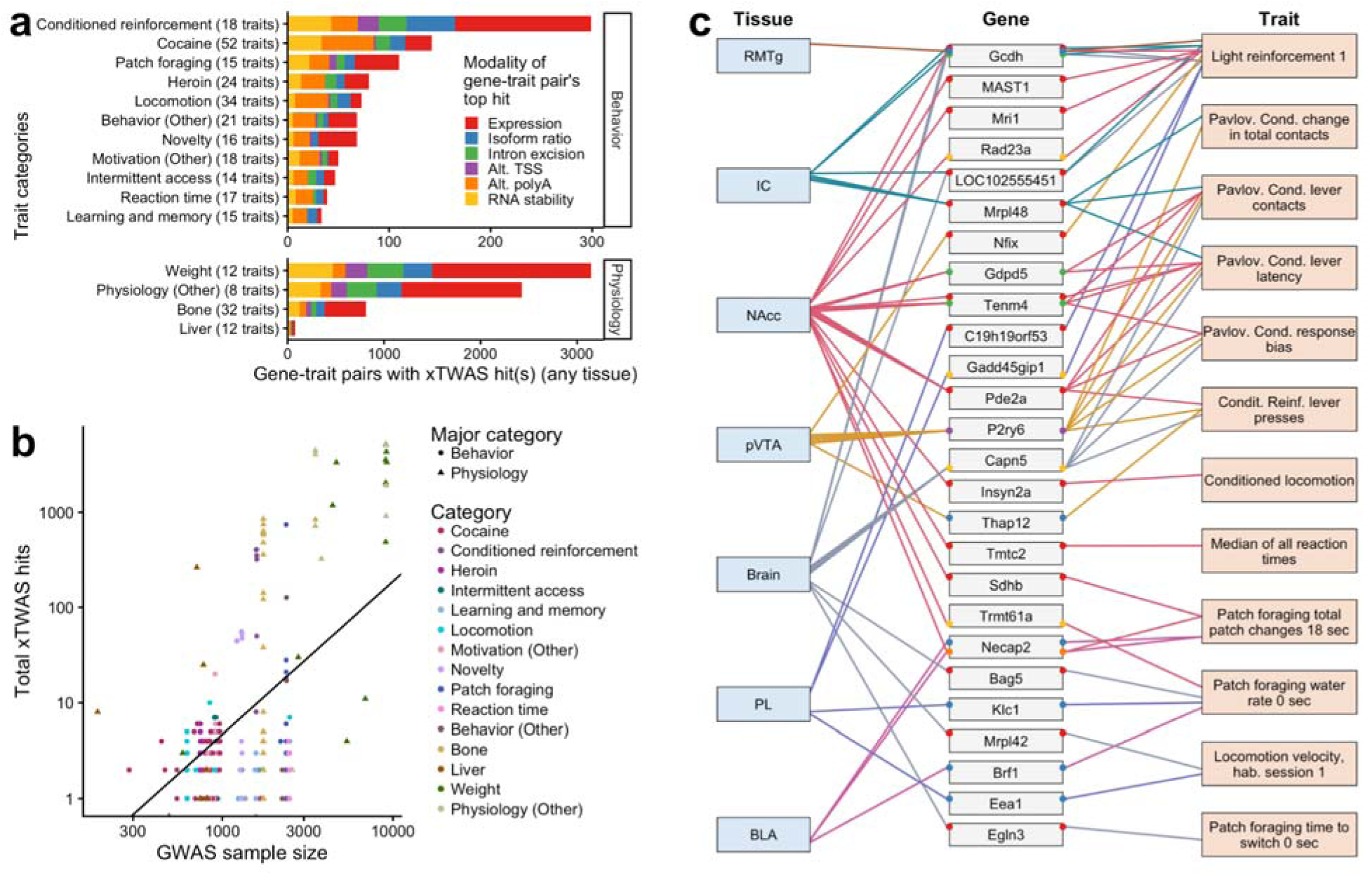
xTWAS associations. **a** Number of unique gene-trait pairs with significant TWAS associations in any tissue and for any transcriptional modality. Gene-trait pairs are grouped by trait category and colored by modality of the transcriptional phenotype with the lowest TWAS p-value for that gene-trait pair. **b** Scatter plot of total xTWAS hits (log scale) versus GWAS sample size (log scale) for each trait. Points are colored by trait category. The ordinary least squares regression line, fitted to log-transformed values, is overlaid. **c** All colocalizing xTWAS hits for brain tissues and behavioral traits. The tissue-gene and gene-trait lines representing each hit are colored by the tissue. Dots at the gene connections are colored by the hit’s modality.

Looking specifically at hits with strong colocalization evidence, for behavioral traits, and in brain tissues, there were 55 hits involving 26 genes and 12 traits in seven tissues (**Figure 4c**). For example, *Egln3* expression in whole brain was associated with “Patch foraging time to switch 0 sec” (**Figure 5a**), and *Insyn2a* expression in nucleus accumbens core was associated with “Conditioned locomotion” (**Figure 5b**).

**Figure 5.**
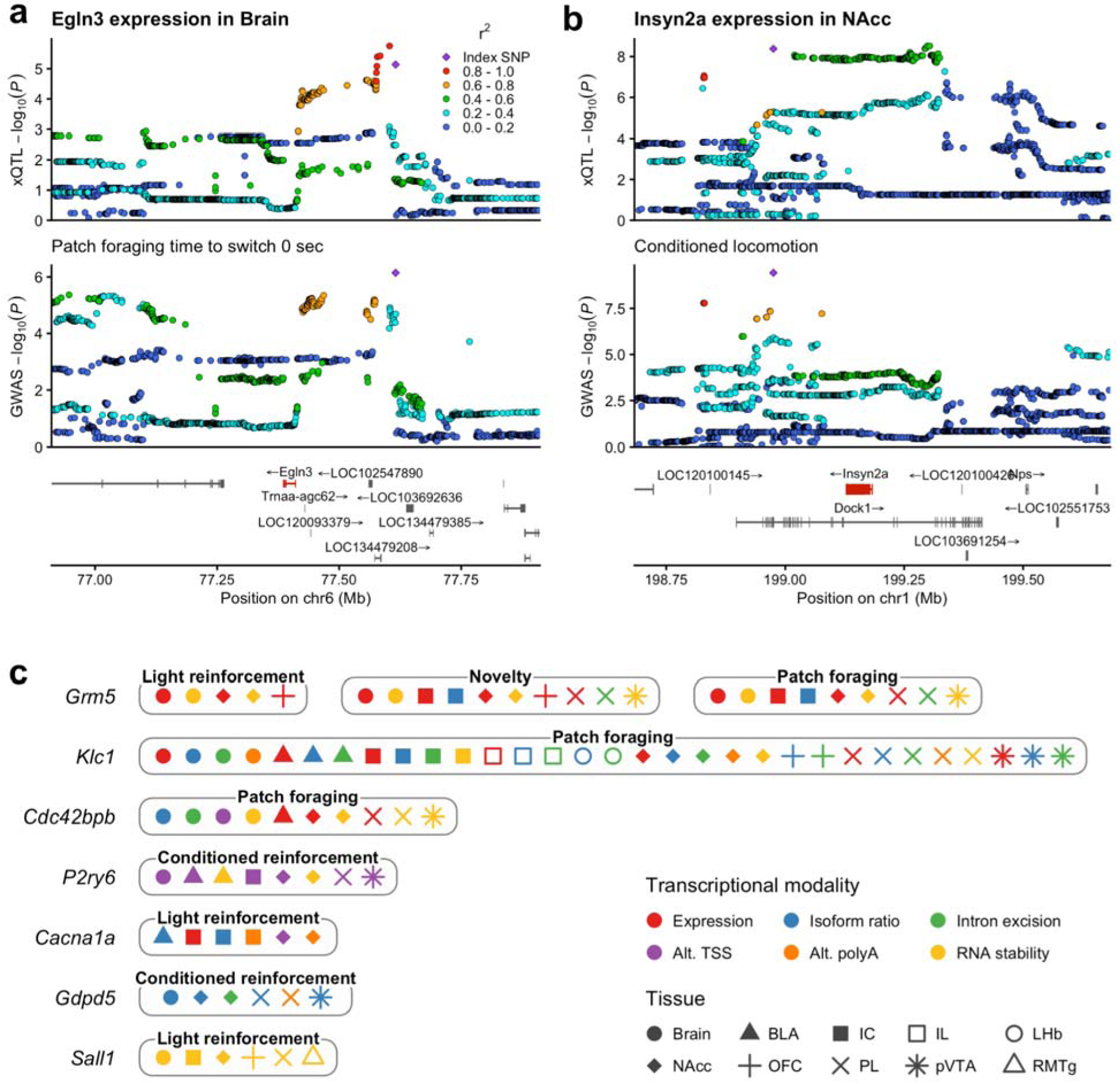
Two examples of colocalizing xTWAS associations. **a** Locuszoom plots for variant associations with *Egln3* expression in whole brain tissue and the “Patch foraging time to switch 0 sec” trait. **b** Locuszoom plots for variant associations with *Insyn2a* expression in nucleus accumbens core and the “Conditioned locomotion” trait. Plot windows include 500 kb on either side of the gene’s transcription start site. **c** xTWAS hits for behavioral traits in brain tissues for a selection of brain-related genes. One symbol is shown per unique gene, trait category, transcriptional modality, tissue combination.

A number of behavior-related genes had associations, though not necessarily with strong colocalization evidence, in brain tissue and for behavioral traits (**Figure 5c**, **Table 1**).

**Table 1.** Selection of brain-related genes with multiple xTWAS hits. Only xTWAS hits for behavioral traits and in brain tissues are counted here.

| **Gene ID** | **Functional relevance** | **xTWAS hit traits** | **xTWAS hit tissues** | **xTWAS hit modalities** |
|---|---|---|---|---|
| *Grm5* | *Grm5* (glutamate metabotropic receptor 5) levels in rats have been shown to affect behaviors related to autism spectrum disorder<sup>23</sup>. | 9 | 6 | 4 |
| *Klc1* | Klc1 (kinesin light chain 1) is crucial for axonal transport, and its reduction in mice leads to axonal defects and increased amyloid deposition<sup>24</sup>. Alternative splicing of *Klc1* contributes to Alzheimer's Disease<sup>25</sup>. | 1 | 9 | 5 |
| *Cdc42bpb* | Deleterious variants in human *CDC42BPB* (Cdc42-binding protein kinase beta) have been associated with neurodevelopmental disorders<sup>26</sup>. | 1 | 5 | 5 |
| *P2ry6* | *P2ry6* (pyrimidinergic receptor P2Y6) codes for a receptor on the surface of microglia and has been shown to mediate aging-induced loss of synapses and memory in mice<sup>27</sup>. | 5 | 6 | 2 |
| *Gdpd5* | Loss of *GDPD5* (glycerophosphodiester phosphodiesterase 2) in mice leads to neurodegenerative changes and memory impairment<sup>28</sup>. | 4 | 4 | 3 |
| *Sall1* | SALL1 (spalt like transcription factor 1), a transcriptional regulator, is important for microglial identity, and its inactivation alters neurogenesis<sup>29</sup>. | 1 | 6 | 1 |
| *Cacna1a* | *Cacna1a* (calcium voltage-gated channel subunit alpha1) codes for a neuronal calcium channel subunit, and its alteration in humans leads to paroxysmal movement disorders<sup>30</sup>. | 1 | 3 | 4 |

Additionally, the *Ccny* gene had TWAS hits for 40 traits, 37 of which were behavioral traits and all of which involved expression in the pVTA brain region. CCNY has been found to modulate spatial learning and memory flexibility in mice^21^, and to inhibit long-term potentiation of synaptic strength in rat neurons^22^.

We tested all xTWAS hits for enrichment in gene sets for ontologies, pathways, and protein domains (see **Methods**). Focusing on the behavioral traits, we found significant enrichment at false discovery rate (FDR) ≤10% of gene sets in xTWAS hits for multiple trait categories, with some enrichments detected for xTWAS hits of specific modalities (**Supplementary Figure S6**). Due to the complex biology of behavior and its regulation, these gene sets are not expected to have consistent strong enrichments in specific molecular concepts. Nevertheless, relevant enrichments were observed, such as “sensory perception of sound” and “sensory perception of mechanical stimulus” as the top two enriched sets for reaction time xTWAS hits, led by two genes, *Alms1* and *Clrn1*. *Alms1* isoform ratios were associated with multiple reaction time traits in the rostromedial tegmental nucleus (RMTg) brain region, while *Clrn1* expression was positively associated with two traits representing reaction time in the basolateral amygdala (BLA) brain region.

## Discussion

In this study, we introduced a large collection of RNA-seq samples from several brain regions and other tissues of outbred rats, deeply quantified their transcriptomes, and performed multimodal genetic analyses. We mapped cis-xQTLs, demonstrated their application to transcriptomic prediction for genes of interest, and identified transcriptional phenotypes genetically associated with a variety of complex traits assembled from a set of outbred rat GWAS studies.

We show that genetic regulation affects multiple modalities of transcriptional variation, highlighting the importance of comprehensive transcriptomic profiling for identifying potential molecular mediators of complex traits. Expression levels yielded the largest number of heritable phenotypes, leading to the most xQTLs, transcriptional phenotype predictions, and xTWAS associations. However, this is not a fixed biological truth, but rather probably reflects the fact that expression uses a larger number of reads and is thus measured more accurately.

Improvements in methods, gene annotations, and sample sizes, along with quantification of additional modalities, will deepen and expand the captured heritable transcriptional variation.

We showed that samples of the same tissue from multiple cohorts, which differed in HS generation number, sequencing technology, and other potential batch effects, can be analyzed as one high-powered dataset. This allows for an ongoing accumulation of datasets to build on existing genetic mapping power. The cross-tissue mash eQTL analysis further demonstrated that different tissue datasets collected from outbred rats can combine information to detect more eQTLs in each tissue.

TWAS analysis provides candidate gene mediators of complex traits, but these associations do not directly implicate the gene in a causal role, even when the gene and trait genetic signals strongly colocalize. As discussed previously^15^, the variant could affect the trait through other causal pathways. Experimental follow-ups could determine which candidates are true mediators.

RatGTEx is a valuable resource that is related to other existing efforts. Our study design and online portal have been heavily influenced by the (human) Genotype-Tissue Expression (GTEx) project^31^ (https://gtexportal.org/). Among non-human model organisms it is conceptually similar to GeneNetwork (https://genenetwork.org/; formerly WebQTL^32^) which provides gene expression and other phenotypic data from multiple model organisms, including early versions of HS rat eQTL data. The Rat Genome Database^33^ (https://rgd.mcw.edu/), QTLViewer^34^ (https://qtlviewer.jax.org/), and PhenoGen^35^ (https://phenogen.org/) provide molecular and/or complex trait QTLs in rodents. Another related resource is FarmGTEx (https://www.farmgtex.org/) which provides similar data for various commercially significant farm animals^36^. Several similar resources also exist for plants, including AraQTL^37^ (https://www.bioinformatics.nl/AraQTL/). While each of these has various strengths and weaknesses, currently, RatGTEx is the only such portal that employs Pantry to explore multiple transcriptional modalities.

As reported previously, cis-genetic associations in heterogeneous stock rats tend to have lower resolution than those mapped in humans due to the outbred rat population’s LD structure. While regulatory variation can be mapped sufficiently to generate useful transcriptional predictions and candidate GWAS mediator genes, transcriptional fine-mapping ability is limited. Nevertheless, GWAS studies in outbred rats enable the study of the genetics, and, as demonstrated here, transcriptomic mediation, of many traits that are not experimentally tested in humans.

## Conclusions

This study reports the largest collection to date of transcriptional phenotypes, molecular QTLs, prediction models, and gene-trait associations in *Rattus norvegicus*. These data will be useful for genetic studies in rats for a deeper understanding of the molecular drivers of genetic associations and to help connect those findings to human biology.

## Methods

### Transcriptomic data collection

We collected RNA-seq datasets from eight studies (**Supplementary Table S1**), three of which have been published^18,38,39^, and either provided the primary data analysis of the study or reanalyzed the data. Each study provided data from a cohort of HS rats with almost no overlap of individuals between cohorts. One to five tissues were collected from each cohort, with samples obtained from largely but not entirely overlapping subsets of individuals per tissue within each cohort.

For the whole brain hemisphere tissue dataset, RNA was extracted from frozen tissue microdissections using a TRIzol-based protocol after mechanical homogenization with a bullet blender to maximize tissue disruption and RNA yield. RNA-seq libraries were constructed using the NEBNext® Ultra™ II Directional RNA Library Prep Kit for Illumina following poly(A) enrichment with the NEBNext Poly(A) mRNA Magnetic Isolation Module. RNA-seq libraries were sequenced on the Illumina NovaSeq 6000 sequencer at the IGM core at UC San Diego.

For the PL-NAcc-pVTA dataset, brain regions were dissected from naive HS rats and RNA extraction was conducted using the same methods as those described previously^18^.

For the IC-NAcc-pVTA dataset, animals were tested under intermittent access or long access cocaine self-administration. Whole brains were flash-frozen and stored at -80°C until sectioning. Brains were equilibrated in a cryostat at -10 to -15°C, embedded in O.C.T., and cut coronally onto microscope slides (typically 120 µm sections; pVTA 45 µm). Slides were kept at -20°C after cutting unless dissected immediately. Regions of interest (ROIs) were microdissected on-slide on a prechilled cold block (about -12 to -14°C). Tools were cleaned with 70% ethanol between brain regions. Each ROI was briefly treated with RNALater-ICE prior to cutting. Dissected tissue was transferred into pre-labeled 1.5 mL tubes containing RLT lysis buffer (RNeasy) plus beta-mercaptoethanol, vortexed for at least 30 seconds, and either processed immediately for RNA extraction or stored at -80°C until shipment to UC San Diego. Total RNA was extracted from brain tissue using RNeasy Mini (Qiagen). The sequencing libraries were prepared using KAPA Stranded mRNA-Seq (Roche). RNA-seq libraries were sequenced on the Illumina NovaSeq X Plus sequencer at the Institute for Genomic Medicine (IGM) core at UC San Diego.

### Transcriptomic data processing

The data in RatGTEx and reported here may differ from the results reported in published source studies for two reasons. First, different processing decisions may have been made for each study, while the data in RatGTEx are processed uniformly to facilitate comparison across tissues. Second, the published study results are immutable, while new versions of data in the portal may be released, for example to improve quality or to use updated reference genome and annotations.

We mapped RNA-seq reads to the GRCr8 genome assembly (GCA_036323735.1) using STAR v2.7.11b^40^, and used GRCr8 RefSeq gene annotations for all transcriptomic analysis. We used allele counts from the RNA-seq data to ensure samples were matched to the same individual’s genotypes, as previously described^18^.

We applied Pantry^12^ to each tissue dataset to generate six modalities of transcriptional phenotypes. Samples from the same tissue from different studies were grouped as a single set of samples and analyzed as one dataset. This applied to NAcc, PL, and pVTA brain regions.

TSS and poly(A) site usage was quantified using txrevise^41^ and kallisto^42^. Expression levels and isoform ratios were quantified using kallisto. Intron excision ratios were quantified using RegTools^43^ and LeafCutter^44^. RNA stability was quantified using featureCounts^45^ to count exonic and intronic reads, using their ratio as a model of mRNA stability^46^. For downstream genetic analyses, phenotypes for each modality in each tissue were filtered and normalized. First, those with more than 50% zeros across samples were removed. Then, quantile normalization was performed to align per-sample distributions, followed by per-phenotype inverse-normal transformation.

### Genotypes

Genotypes were generated primarily using low-coverage whole-genome sequencing (approximately 0.25x coverage), with a subset of earlier samples assayed by genotyping-by-sequencing. Reads were aligned to the GRCr8 rat reference genome using BWA-MEM, and genotypes were imputed using STITCH^47^ with a reference panel constructed from whole-genome sequencing of the eight HS founder strains. Repository quality control excluded variants with imputation INFO scores <0.9 or >5% non-Mendelian genotypes, and samples with >5% non-Mendelian genotypes, >10% missing genotypes, or heterozygosity more than five standard deviations from the population mean.

### xQTL mapping

We performed cis-QTL mapping on transcriptional phenotypes using tensorQTL^19^, following the default Pantry Pheast pipeline. This tested variants within 1 Mb of the gene’s TSS, computed permutation-based P-values, and used a 5% FDR threshold across genes to determine significant xQTLs. Stepwise regression was then used to identify any additional, conditionally independent cis-xQTLs for each gene. We used 25 data-driven covariates, including the first five principal components from the LD-pruned genotype matrix and the first 20 principal components from the transcriptional phenotype matrix. We used cross-modality mapping so that the xQTLs reported in this study are conditionally independent across modalities for each gene, keeping up to 10 xQTLs per gene per tissue.

To jointly estimate cis-eQTL effects across tissues, we applied multivariate adaptive shrinkage (mash)^20^, implemented in the R package mashr, to the marginal effect estimates and standard errors from the tissue-specific nominal association scans. Gene-variant pairs were aligned across the 13 tissues, retaining only pairs tested in every tissue. A random sample of 200,000 pairs was used to estimate cross-tissue correlations in estimation error and fit the mixture weights. Data-driven covariance matrices were learned from the union of the most strongly associated variant for each gene in each tissue, selected by nominal P-value, using five principal components followed by extreme deconvolution. These matrices were combined with canonical covariance matrices to model shared and tissue-specific effects. The fitted model was then applied to all retained pairs with mixture weights fixed, yielding posterior effect estimates and local false sign rates (LFSRs). For each gene and tissue, we selected the variant with the lowest LFSR and called the gene an eGene at LFSR ≤ 0.05. Comparisons with individual-tissue mapping were restricted to protein-coding and long noncoding RNA genes represented in the mash analysis.

### GWAS data

We collected GWAS summary statistics for 308 traits from 14 studies on HS rats (**Supplementary Table S2**, **Supplementary Table S5**)^39,48–54^ . Each trait was assigned one or more descriptive tags, and for the purposes of summarization and visualization, each trait was assigned to one category using this process:

1. Remove tags that appear <12 times.
2. For each trait with multiple tags, choose the tag that appears the fewest times.
3. Identify tags where some of its traits are still assigned that tag, but most have been assigned other tags after step 2. For the traits still assigned that tag, use the modified label “{tag} (Other)”.

### xTWAS

We used the Pantry Pheast module to generate TWAS transcriptomic models for each phenotype using FUSION^16^ on the phenotype tables, genotypes, and the same covariates used for xQTL mapping. We used the whole brain hemisphere (Brain) tissue dataset as the LD reference panel. Only models with heritability P-value <0.01 were kept. We then tested TWAS associations using FUSION with default parameters. We used a Bonferroni P-value threshold of 0.05 / 259,088 = 1.93×10^−7^, where 259,088 is the total number of TWAS transcriptional models across tissues and modalities, to determine significant xTWAS hits. Colocalization tests as implemented in the coloc R package^55^ were applied to each TWAS hit, and hits with COLOC.PP4 (posterior probability of shared causal variant) > 0.8 were considered colocalizing TWAS hits for the purposes of this study.

We performed gene-set over-representation analysis on genes implicated by xTWAS. Within each trait category, we pooled hit genes across traits and tissues, separately for each transcriptional modality and across all modalities (“Combined”), counting each gene once per group. Groups containing at least 10 unique hit genes were analyzed. Rat gene sets were obtained from the Bader Lab GeneSetDB collection (GO_K_NCI_BIOC_PF_Rn_eg.GMT, 2010 release), comprising Gene Ontology biological process, cellular component, and molecular function terms; KEGG, NCI, and BioCarta pathways; and Pfam protein domains. Entrez Gene identifiers were mapped to the study’s gene identifiers using the reference genome annotation, supplemented by Rat Genome Database mappings^33^; ambiguous mappings were excluded.

Background universes comprised unique genes represented in the xTWAS model catalog for the corresponding modality, or the union across modalities for Combined analyses. Gene sets were intersected with the relevant background universe, and sets containing 10–500 background genes were retained. Enrichment was assessed using one-sided hypergeometric tests. P values were adjusted using the Benjamini–Hochberg procedure separately within each trait-category–modality group, across retained gene sets containing at least one hit gene.

Enrichments with an adjusted P value ≤0.10 were considered significant.

### Transcriptional phenotype prediction

We used the same transcriptomic models generated for TWAS using FUSION to predict the phenotypes for genotyped rats. We converted the models to individual-level predictors using FUSION’s make_score.R script, and applied them to the genotypes of 24,722 HS rats, a collection comprising the RNA-seq and GWAS cohorts in this study and others, using PLINK2^56^ with the --score parameter. The resulting predictions were Z-scores representing relative deviation from average. Cross-validation accuracy was assessed as part of the original model-fitting stage and compiled using the FUSION.profile_wgt.R script.

## Code availability

Code for running the pipeline from raw data to xQTLs and TWAS models is available at https://github.com/daniel-munro/ratgtex-pipeline. Code for processing data further into files for download and the website’s data application programming interface (API) is available at https://github.com/daniel-munro/ratgtex-server-data. Code for the website itself, including the Flask data API server, is available at https://github.com/daniel-munro/ratgtex. Code for the Rat TWAS Hub is available at https://github.com/daniel-munro/rat-twas-hub. Code for the figures, tables, and analysis in this paper is available at https://github.com/daniel-munro/ratgtex-paper.

## Declarations

### Ethics approval and consent to participate

Animal procedures associated with the generation of the previously unpublished datasets reported in this study were approved by the Institutional Animal Care and Use Committee (IACUC) at UC San Diego, The University of Tennessee Health Science Center, Medical College of Wisconsin, and University at Buffalo. Consent to participate was not applicable.

### Consent for publication

Not applicable

### Availability of data and materials

Data presented in this study correspond to RatGTEx (RRID:SCR_022145) v4. Transcriptomic and genomic data including transcriptional phenotypes, genotypes, xQTLs, and metadata are available at https://ratgtex.org/download/. xTWAS results can be downloaded from https://twas.ratgtex.org/traits/. Raw RNA-seq data have been deposited at GEO accessions GSE173141, GSE196572, GSE200453, GSE201236, GSE319591, GSE325483, GSE325609, GSE325289, and GSE325599.

### Authors’ contributions

D.B.M. processed, analyzed, and visualized the data, implemented the web resources, and wrote the manuscript. A.A.P. and P.M. secured funding, planned and supervised the analyses, and edited the manuscript. H.C., L.C.S.W., S.H.M., P.J.M., F.T., T.C.J., M.M.J., and R.H. produced the RNA-seq datasets and provided feedback for the RatGTEx resource. L.M.S. provided analysis and feedback for the RatGTEx resource. K.-M.H.N., C.L.S.P., H.B., K.A.C., Y.W., R.W., and K.H. handled tissue samples and library preparation. D.C., B.B.J., G.A., M.M., and E.K. performed genotyping and curated sequencing data. O.P. coordinated sample and data collection. A.G. planned and interpreted the results of the TWAS analyses. C.K., J.A.T., B.N.K., and B.M.T. performed experimental design, data collection, and analysis for complex traits. P.J.M., T.C.J., M.M.J., S.F., D.M.D., S.H.M., D.J.A., O.G., P.W.K., C.V., D.D., and A.L. produced phenotypes for GWAS. All authors read and approved the final manuscript.

## Acknowledgements

This publication includes data generated at the UC San Diego IGM Genomics Center utilizing an Illumina NovaSeq X Plus that was purchased with funding from a National Institutes of Health Shared Instrumentation Grant (#S10 OD026929). RatGTEx is produced by the NIDA center for genetic studies of drug abuse in outbred rats (P50DA037844) and is not a project of the GTEx Consortium. Code for the data processing pipeline and portal visualizations released by the GTEx Consortium was adapted for RatGTEx.

## References

1. Ren, Y. & Palmer, A. A. Behavioral Genetic Studies in Rats. in Rat Genomics (eds Hayman, G. T., Smith, J. R., Dwinell, M. R. & Shimoyama, M.) 319–326 (Springer, New York, NY, 2019). doi:10.1007/978-1-4939-9581-3_16.

2. Aitman, T. J. et al. Progress and prospects in rat genetics: a community view. Nat. Genet. 40, 516–522 (2008).

3. Parker, C. C. et al. Rats are the smart choice: Rationale for a renewed focus on rats in behavioral genetics. Neuropharmacology 76, 250–258 (2014).

4. Solberg Woods, L. C. QTL mapping in outbred populations: successes and challenges. Physiol. Genomics 46, 81–90 (2014).

5. Moreno-Moral, A. & Petretto, E. From integrative genomics to systems genetics in the rat to link genotypes to phenotypes. Dis. Model. Mech. 9, 1097–1110 (2016).

6. Ellenbroek, B. & Youn, J. Rodent models in neuroscience research: is it a rat race? Dis. Model. Mech. 9, 1079–1087 (2016).

7. Gileta, A. F. et al. Genetic characterization of outbred Sprague Dawley rats and utility for genome-wide association studies. PLOS Genet. 18, e1010234 (2022).

8. Solberg Woods, L. C. & Palmer, A. A. Using Heterogeneous Stocks for Fine-Mapping Genetically Complex Traits. in Rat Genomics (eds Hayman, G. T., Smith, J. R., Dwinell, M. R. & Shimoyama, M.) 233–247 (Springer, New York, NY, 2019). doi:10.1007/978-1-4939-9581-3_11.

9. Baud, A. et al. Combined sequence-based and genetic mapping analysis of complex traits in outbred rats. Nat. Genet. 45, 767–775 (2013).

10. Woods, L. C. S. & Mott, R. Heterogeneous Stock Populations for Analysis of Complex Traits. in Systems Genetics: Methods and Protocols (eds Schughart, K. & Williams, R. W.) 31–44 (Springer, New York, NY, 2017). doi:10.1007/978-1-4939-6427-7_2.

11. Lappalainen, T., Li, Y. I., Ramachandran, S. & Gusev, A. Genetic and molecular architecture of complex traits. Cell 187, 1059–1075 (2024).

12. Munro, D. et al. Multimodal analysis of RNA sequencing data powers discovery of complex trait genetics. Nat. Commun. 15, 10387 (2024).

13. Zhou, J. L. et al. Single-nucleus genomics in outbred rats with divergent cocaine addiction-like behaviors reveals changes in amygdala GABAergic inhibition. Nat. Neurosci. 26, 1868–1879 (2023).

14. Campo, P. et al. Ivermectin reduces withdrawal-induced alcohol intake in rats: Association with CeA GABAergic enhancement and *P2rx4* genetic liability. Neuropharmacology 289, 110881 (2026).

15. Gamazon, E. R. et al. A gene-based association method for mapping traits using reference transcriptome data. Nat. Genet. 47, 1091–1098 (2015).

16. Gusev, A. et al. Integrative approaches for large-scale transcriptome-wide association studies. Nat. Genet. 48, 245–252 (2016).

17. Santhanam, N. et al. RatXcan: A framework for cross-species integration of genome-wide association and gene expression data. PLOS Genet. 21, e1011583 (2025).

18. Munro, D. et al. The regulatory landscape of multiple brain regions in outbred heterogeneous stock rats. Nucleic Acids Res. gkac912 (2022) doi:10.1093/nar/gkac912.

19. Taylor-Weiner, A. et al. Scaling computational genomics to millions of individuals with GPUs. Genome Biol. 20, 228 (2019).

20. Urbut, S. M., Wang, G., Carbonetto, P. & Stephens, M. Flexible statistical methods for estimating and testing effects in genomic studies with multiple conditions. Nat. Genet. 51, 187–195 (2019).

21. Seo, J. et al. Cyclin Y regulates spatial learning and memory flexibility through distinct control of the actin pathway. Mol. Psychiatry 28, 1351–1364 (2023).

22. Cho, E. et al. Cyclin Y inhibits plasticity-induced AMPA receptor exocytosis and LTP. Sci. Rep. 5, 12624 (2015).

23. Eshraghi, A. A. et al. Genetic ablation of metabotropic glutamate receptor 5 in rats results in an autism-like behavioral phenotype. PLOS ONE 17, e0275937 (2022).

24. Stokin, G. B. et al. Axonopathy and Transport Deficits Early in the Pathogenesis of Alzheimer’s Disease. Science 307, 1282–1288 (2005).

25. Morihara, T., et al. Transcriptome analysis of distinct mouse strains reveals kinesin light chain-1 splicing as an amyloid-β accumulation modifier. Proc. Natl. Acad. Sci. 111, 2638–2643 (2014).

26. Chilton, I. et al. De novo heterozygous missense and loss-of-function variants in CDC42BPB are associated with a neurodevelopmental phenotype. Am. J. Med. Genet. A. 182, 962–973 (2020).

27. Dundee, J. M., Puigdellívol, M., Butler, R., Cockram, T. O. J. & Brown, G. C. P2Y6 receptor-dependent microglial phagocytosis of synapses mediates synaptic and memory loss in aging. Aging Cell 22, e13761 (2023).

28. Daudelin, D. et al. Loss of GDE2 leads to complex behavioral changes including memory impairment. Behav. Brain Funct. 20, 7 (2024).

29. Buttgereit, A. et al. Sall1 is a transcriptional regulator defining microglia identity and function. Nat. Immunol. 17, 1397–1406 (2016).

30. Indelicato, E. & Boesch, S. From Genotype to Phenotype: Expanding the Clinical Spectrum of CACNA1A Variants in the Era of Next Generation Sequencing. Front. Neurol. 12, (2021).

31. The GTEx Consortium. The GTEx Consortium atlas of genetic regulatory effects across human tissues. Science 369, 1318–1330 (2020).

32. Wang, J., Williams, R. W. & Manly, K. F. WebQTL. Neuroinformatics 1, 299–308 (2003).

33. Vedi, M. et al. 2022 updates to the Rat Genome Database: a Findable, Accessible, Interoperable, and Reusable (FAIR) resource. Genetics 224, iyad042 (2023).

34. Vincent, M., et al. QTLViewer: an interactive webtool for genetic analysis in the Collaborative Cross and Diversity Outbred mouse populations. G3 GenesGenomesGenetics 12, jkac146 (2022).

35. Bennett, B. et al. Genetical Genomic Analysis of Complex Phenotypes Using the PhenoGen Website. Behav. Genet. 41, 625–628 (2011).

36. Fang, L. et al. The Farm Animal Genotype–Tissue Expression (FarmGTEx) Project. Nat. Genet. 57, 786–796 (2025).

37. Nijveen, H. et al. AraQTL – workbench and archive for systems genetics in Arabidopsis thaliana. Plant J. 89, 1225–1235 (2017).

38. Hong-Le, T. et al. Genetic Mapping of Multiple Traits Identifies Novel Genes for Adiposity, Lipids, and Insulin Secretory Capacity in Outbred Rats. Diabetes 72, 135–148 (2022).

39. Fowler, S. et al. Genome-wide association study finds multiple loci associated with intraocular pressure in HS rats. Front. Genet. 13, (2023).

40. Dobin, A. et al. STAR: ultrafast universal RNA-seq aligner. Bioinformatics 29, 15–21 (2013).

41. Alasoo, K. et al. Genetic effects on promoter usage are highly context-specific and contribute to complex traits. eLife 8, e41673 (2019).

42. Bray, N. L., Pimentel, H., Melsted, P. & Pachter, L. Near-optimal probabilistic RNA-seq quantification. Nat. Biotechnol. 34, 525–527 (2016).

43. Cotto, K. C. et al. Integrated analysis of genomic and transcriptomic data for the discovery of splice-associated variants in cancer. Nat. Commun. 14, 1589 (2023).

44. Li, Y. I. et al. Annotation-free quantification of RNA splicing using LeafCutter. Nat. Genet. 50, 151–158 (2018).

45. Liao, Y., Smyth, G. K. & Shi, W. featureCounts: an efficient general purpose program for assigning sequence reads to genomic features. Bioinformatics 30, 923–930 (2014).

46. Gaidatzis, D., Burger, L., Florescu, M. & Stadler, M. B. Analysis of intronic and exonic reads in RNA-seq data characterizes transcriptional and post-transcriptional regulation. Nat. Biotechnol. 33, 722–729 (2015).

47. Davies, R. W., Flint, J., Myers, S. & Mott, R. Rapid genotype imputation from sequence without reference panels. Nat. Genet. 48, 965–969 (2016).

48. Hughson, A. R. et al. Incentive salience attribution, “sensation-seeking” and “novelty-seeking” are independent traits in a large sample of male and female heterogeneous stock rats. Sci. Rep. 9, 2351 (2019).

49. King, C. P. et al. Sensitivity to food and cocaine cues are independent traits in a large sample of heterogeneous stock rats. Sci. Rep. 11, 2223 (2021).

50. Lara, M. K. et al. Genome-wide association study of delay discounting in Heterogeneous Stock rats. Genes Brain Behav. 23, e12909 (2024).

51. King, C. P. et al. Genetic Loci Influencing Cue-Reactivity in Heterogeneous Stock Rats. Genes Brain Behav. 24, e70018 (2025).

52. Kuhn, B. N. et al. Genome-wide association study reveals multiple loci for nociception and opioid consumption behaviors associated with heroin vulnerability in outbred rats. Mol. Psychiatry 30, 3363–3375 (2025).

53. Lara, M. K. et al. Genome-wide association study of cocaine self-administration behavior in Heterogeneous Stock rats. Nat. Commun. 17, 4876 (2026).

54. Tatom, Z. et al. Genome-wide association study in Heterogeneous Stock rats identifies genetic loci associated with aversion-based learning and cocaine aversion. 2026.08.07.743619 Preprint at 10.64898/2026.08.07.743619 (2026).

55. Giambartolomei, C. et al. Bayesian Test for Colocalisation between Pairs of Genetic Association Studies Using Summary Statistics. PLOS Genet. 10, e1004383 (2014).

56. Chang, C. C. et al. Second-generation PLINK: rising to the challenge of larger and richer datasets. GigaScience 4, s13742–015-0047–8 (2015).

