## Supplementary Materials for "Deep transcriptomic analysis of molecular mediators of complex traits in heterogeneous stock (HS) rats"

### Supplementary Information

**In this file:**
Supplementary Tables S1-S2 and descriptions for S3-S6
Supplementary Figures S1-S6

### Supplementary Tables

**Supplementary Table S1**. The 19 tissue datasets collected for RatGTEx and Rat TWAS Hub and analyzed in this study. Datasets from the same tissue from different studies were merged and analyzed as one dataset. Sample sizes reflect the samples passing quality control filtering and that have genotype data.

| Tissue | Abbreviation | Samples | Originating study |
|---|---|---|---|
| Infralimbic cortex | IL | 83 | Munro et al. 2022 (PMID: 36263809). P50DA037844 Y1-5, Project 2. PI: Hao Chen, University of Tennessee Health Science Center |
| Lateral habenula | LHb | 82 | |
| Nucleus accumbens core | NAcc | 77 | |
| Orbitofrontal cortex | OFC | 82 | |
| Prelimbic cortex | PL | 82 | |
| Visceral adipose | Adipose | 411 | Hong-Le et al. 2022 (PMID: 36219827). R01DK106386. PI: Leah C. Solberg Woods, Wake Forest University |
| Liver | Liver | 411 | |
| Eye | Eye | 53 | Fowler et al. 2023 (PMID: 36793389). R01EY021200. PI: Monica M. Jablonski, University of Tennessee Health Science Center |
| Basolateral amygdala | BLA | 189 | U01DA046077. PI: Suzanne H. Mitchell and Robert Hitzemann, Oregon Health & Science University |
| Nucleus accumbens core | NAcc | 191 | |
| Prelimbic cortex | PL | 192 | |

| | | | |
|---|---|---|---|
| Brain hemisphere | Brain | 342 | PI: Francesca Telese, UC San Diego |
| Nucleus accumbens core | NAcc | 159 | P50DA037844 Y6-10, Project 2. PI: Hao Chen, University of Tennessee Health Science Center |
| Prelimbic cortex | PL | 130 | |
| Posterior ventral tegmental area | pVTA | 152 | |
| Rostromedial tegmental nucleus | RMTg | 92 | U01DA044468. PI: Thomas C. Jhou, University of Maryland |
| Agranular insular cortex | IC | 148 | P50DA037844 Y6-10, Project 1. PI: Paul J. Meyer, University at Buffalo |
| Nucleus accumbens core | NAcc | 143 | |
| Posterior ventral tegmental area | pVTA | 141 | |

**Supplementary Table S2**. The 14 studies from which GWAS summary statistics were collected for use in xTWAS. Publications cover the phenotyping and/or GWAS for that dataset. Studies with no publication listed have not yet been published. The number of traits reflects the number used in Rat TWAS Hub and the present study rather than the full set measured in the original study.

| **Title** | **Principal Investigator** | **Number of traits** | **Publication** |
|---|---|---|---|
| Genetic studies of incentive salience | Shelly Flagel, University of Michigan | 49 | Hughson et al. 2019 (PMID: 30787409) |
| Neurogenetic substrates of cocaine addiction | Paul J. Meyer, University at Buffalo | 42 | |
| Association between behavioral regulation and cocaine cue preference 2 | David M. Dietz, University at Buffalo | 40 | King et al. 2025 (PMID: 40049657) |
| Identification of genes regulating bone matrix composition and quality | Douglas J. Adams, University of Colorado | 31 | |
| Identification of genetic variants that contribute to compulsive cocaine intake in rats | Olivier George, University of California San Diego | 27 | Lara et al. 2026 (PMID: 42277005) |
| The genetic basis of opioid dependence vulnerability in a rodent model | Peter W. Kalivas, Medical University of South Carolina | 24 | Kuhn et al. 2025 (PMID: 40000848) |

| | | | |
|---|---|---|---|
| Identification of genetic features of delay discounting using a heterogeneous stock rat model | Suzanne H. Mitchell, Oregon Health & Science University | 23 | Lara et al. 2024 (PMID: 39119916) |
| Association between behavioral regulation and cocaine cue preference 1 | Paul J. Meyer, University at Buffalo | 21 | King et al. 2021 (PMID: 33500444) |
| Physiological and morphological phenotypes | | 15 | |
| Genomic analysis of avoidance learning in addiction | Thomas C. Jhou, University of Maryland | 12 | Tatom et al. 2026 (preprint: https://doi.org/10.64898/2026.08.07.743619) |
| GWAS of Hepatic Mineral and Trace Element Concentrations in Outbred Heterogeneous Stock Rats | Christopher Vulpe, University of Florida | 12 | |
| GWAS for 24-hour food and fluid intake in heterogeneous stock rats | Derek Daniels, University at Buffalo | 7 | |
| Genetics of muscles in hind limb | Arimantas Lionikas, University of Aberdeen | 4 | |
| Genome-wide association study finds multiple loci associated with intraocular pressure in HS rats | Monica M. Jablonski, University of Tennessee Health Science Center | 1 | Fowler et al. 2023 (PMID: 36793389) |

**Supplementary Table S3**: Counts of transcriptional phenotypes, xQTLs, and xGenes per tissue

**Supplementary Table S4**: Cross-tissue mash eQTL results

**Supplementary Table S5**: TWAS trait information

**Supplementary Table S6**: xTWAS hit and gene counts per trait per tissue

### Supplementary Figures

**Supplementary Figure S1: Factors correlated to number of xQTLs per tissue.**

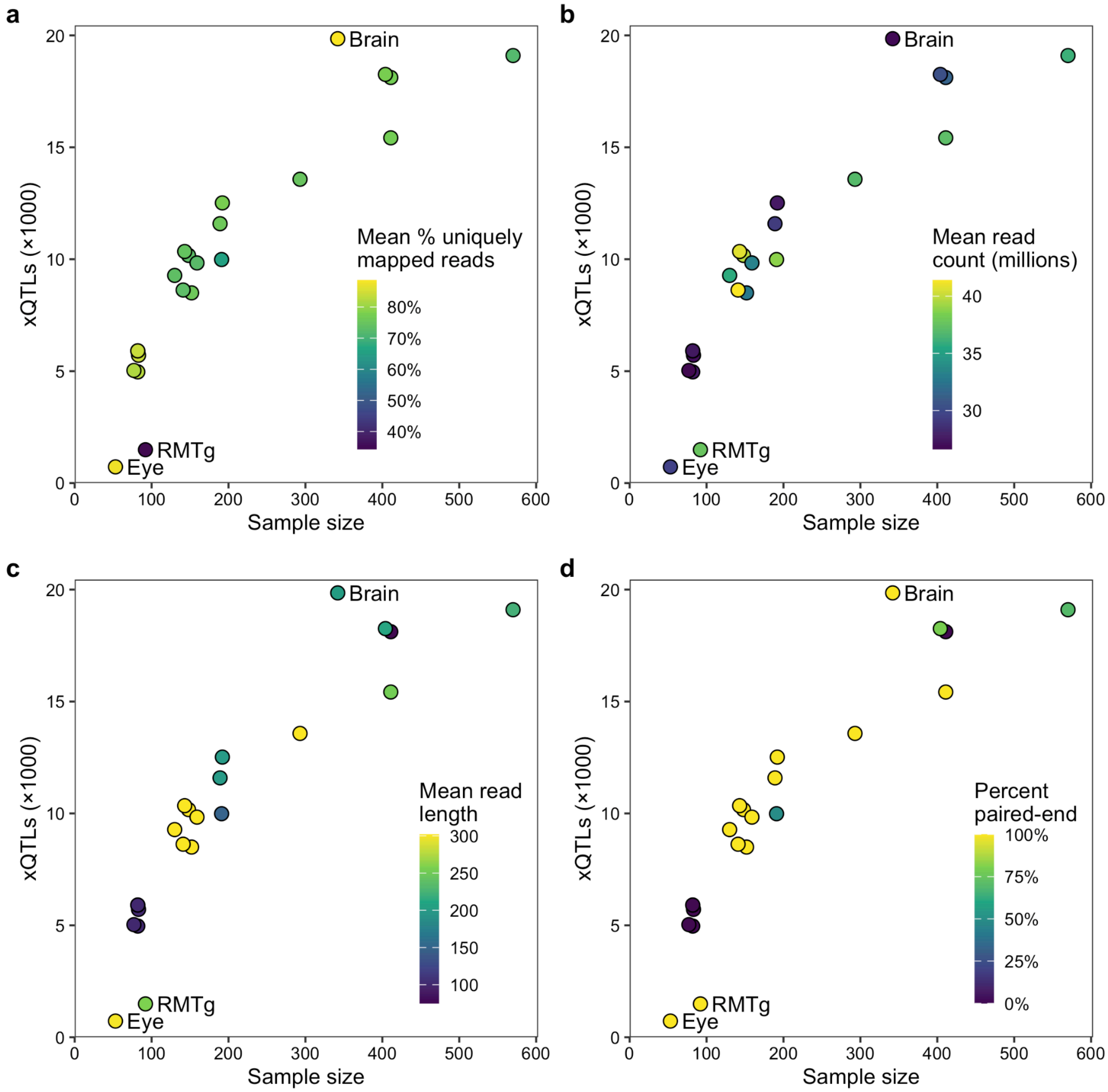


Scatter plots of the number of independent xQTLs versus sample size per tissue. For each tissue with samples from multiple cohorts, the number of xQTLs and samples for each pre-merged sample set and for the merged set are both shown. Each panel shows the same scatter plot, but with points colored by different statistics: tissue's mean percent uniquely mapped reads, mean read count, mean read length (reported as twice the length of each end for paired-end), and percent of samples being paired-end.

**Supplementary Figure S2: Dataset-associated phenotype structure before and after phenotype PC adjustment.**

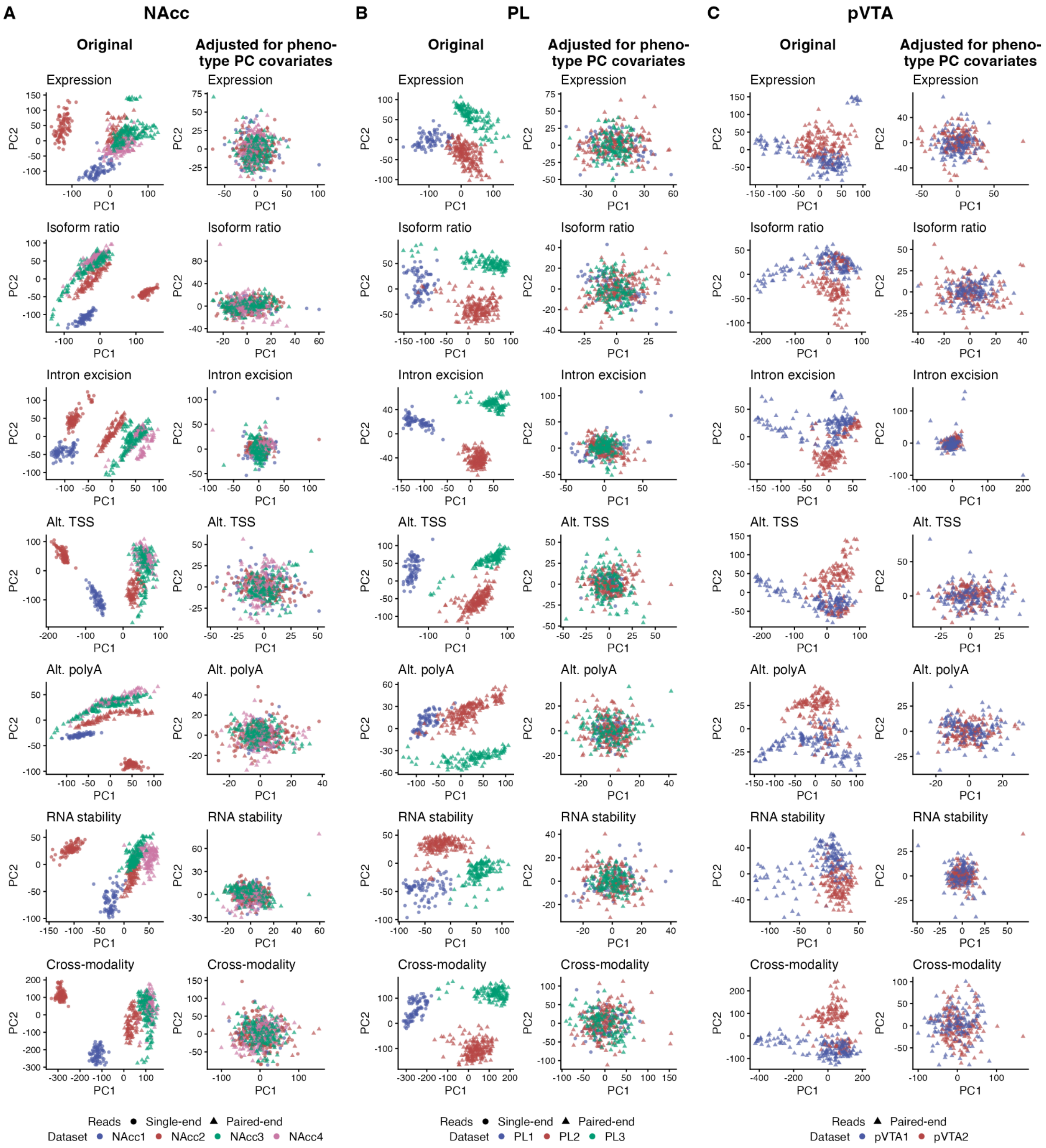


Scores for the first two principal components (PCs) are shown for the three multi-dataset tissues, across six transcriptional modalities and the joint cross-modality analysis. Within each tissue, left columns show PCs of the original normalized phenotypes; right columns show PCs recomputed after residualization on the 20 phenotype PC covariates. Point shapes indicate whether an RNA-seq sample is single-end or paired-end, and color indicates which source dataset the sample came from. Dataset numbers and colors do not necessarily correspond across tissues.

**Supplementary Figure S3: Transcriptional phenotype prediction accuracy across tissues and modalities.**

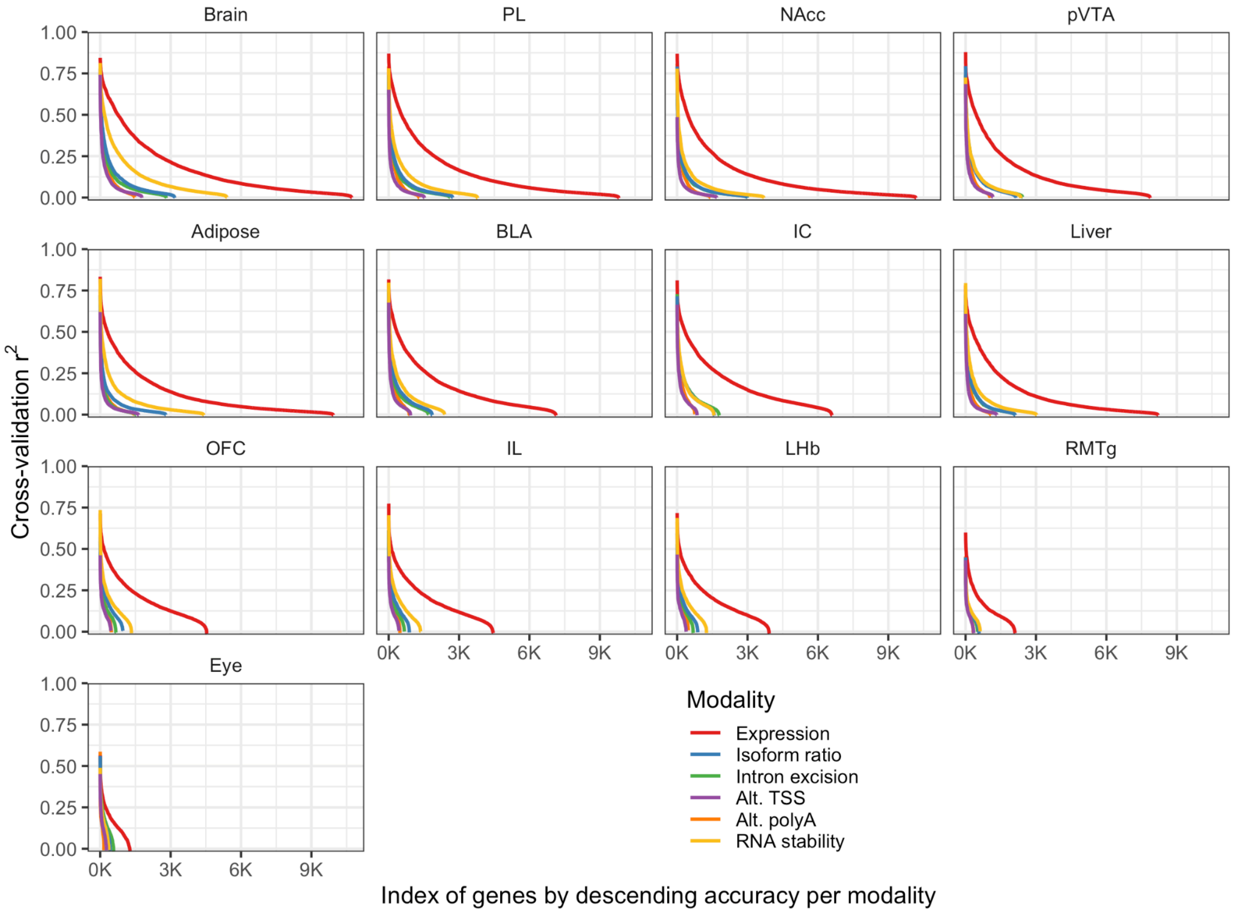


Accuracy for the transcriptional phenotypes with cis-genetic prediction models in each tissue, determined by having heritability P-value <0.01. Within each modality in each tissue, genes were ordered on the x-axis by descending cross-validation adjusted $r^2$, or mean $r^2$ for genes with multiple predicted phenotypes of the same modality.

**Supplementary Figure S4: Colocalizing TWAS hits versus transcriptomic prediction model accuracy**

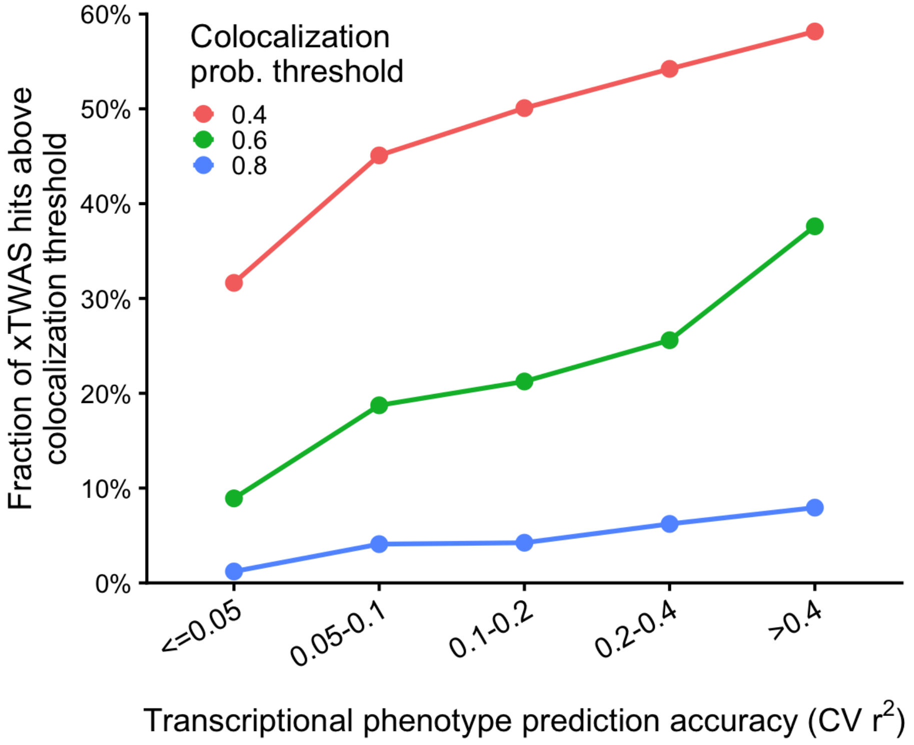


For each range of transcriptional phenotype prediction accuracy $r^2$ values, the fractions of significant xTWAS hits for those phenotypes that colocalize with probability above the indicated thresholds are shown.

**Supplementary Figure S5: Porcupine plots of all xTWAS results**

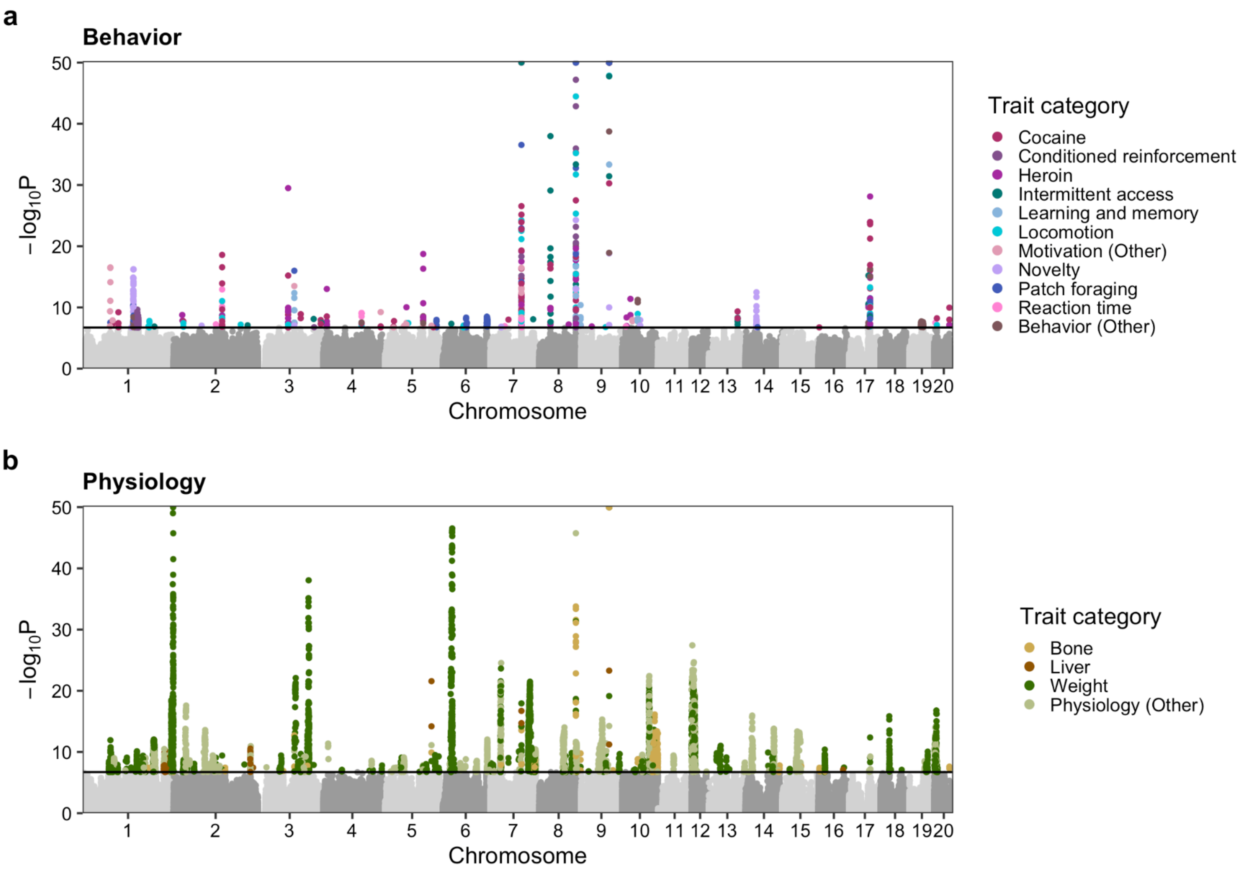


**a** xTWAS associations for all tested behavioral traits. Significant associations (those above the Bonferroni threshold $1.93\times10^{-7}$ shown as a black line) are colored by trait category. Any $-\log_{10}$P-values >=50 are shown as 50. **b** The same plot for all tested physiological traits.

**Supplementary Figure S6:** Gene-set enrichment among xTWAS hit genes for behavioral traits.

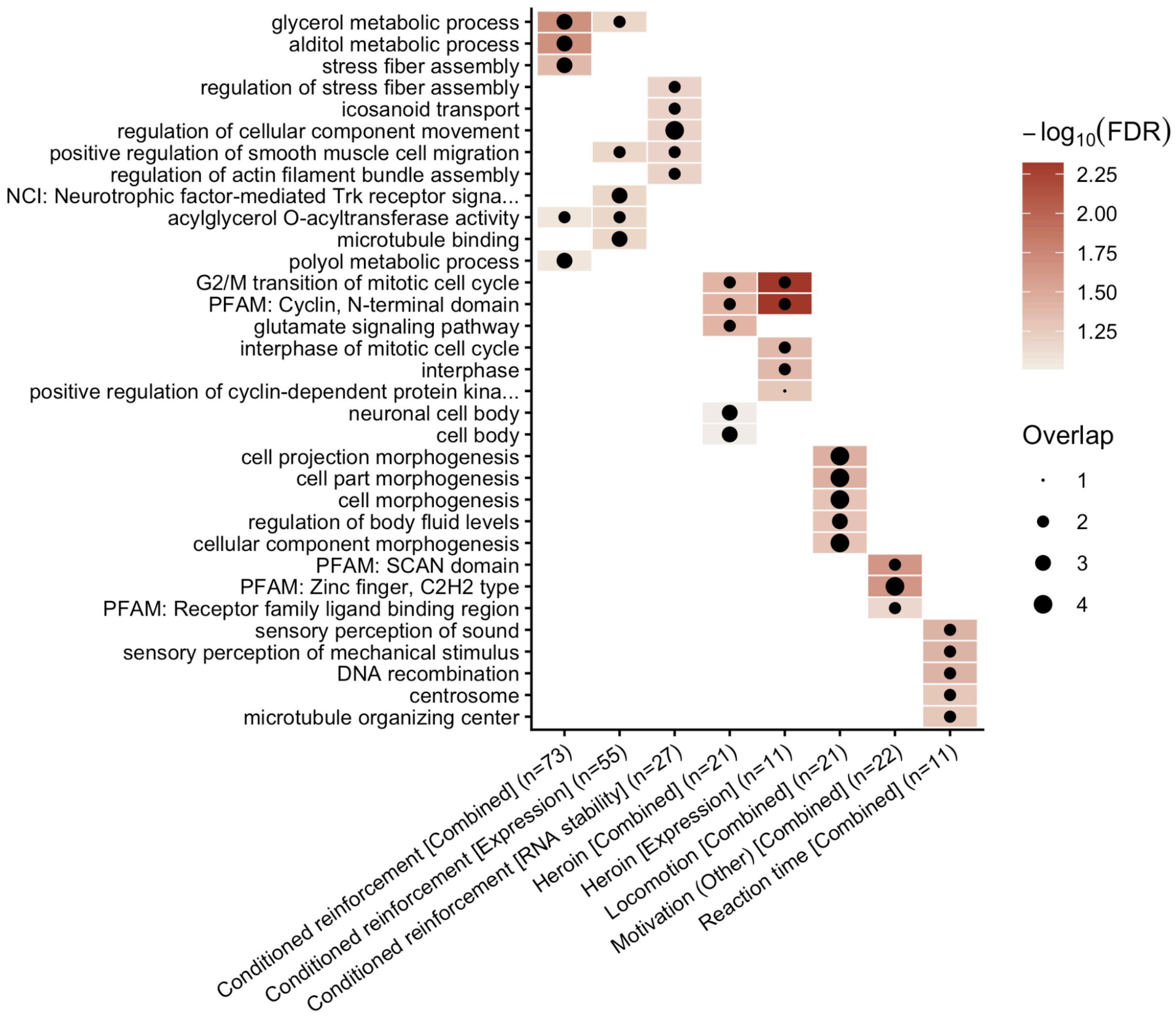


For each behavioral trait category, unique genes with xTWAS hits were pooled across traits and tissues and analyzed separately for each RNA modality and for their union (“Combined”). Up to five significant gene sets (FDR ≤0.10) with the lowest adjusted P-values are displayed per group. Tile color indicates $-\log_{10}(\mathrm{FDR})$, and black point size indicates the number of hit genes overlapping the gene set. Counts in column labels indicate the number of unique hit genes in each group. Rows are sorted by the category of their strongest displayed enrichment, then lowest FDR. Behavioral hit groups with no significant enrichments are not shown.
